# Adipocyte IL-22RA1 signaling promotes structural and functional remodeling of white adipose tissue following acute intestinal damage

**DOI:** 10.1101/2025.11.14.688505

**Authors:** Ankita Singh, Kiyoshi Shiomitsu, Stephen J. Gaudino, Hafiz Tariq, Pawan Kumar

## Abstract

Interleukin (IL)-22 has been shown to play an important role in intestinal host defense and ameliorating high-fat diet (HFD)-induced metabolic disorders primarily through signaling in intestinal epithelial cells. Adipocytes have emerged as key immune–metabolic regulators that influence intestinal inflammation in inflammatory bowel disease (IBD). However, the role of IL-22RA1 signaling in adipocytes has not been explored. In the present study, we examined the role of IL-22RA1 signaling in adipocytes in response to dextran sulfate sodium (DSS)-mediated gut inflammation. To do this, we subjected adipocyte specific *Il22ra1* knockout mice (*Il22ra1^Adipo^*) to intestinal inflammation under normal chow and HFD conditions. Compared to littermate controls, *Il22ra1^Adipo^* mice displayed significant weight loss on day 9 of DSS treatment and showed altered expression of immune response– and lipid metabolism–related genes in white adipose tissue (WAT) under a normal chow diet. Notably, when mice were primed with HFD prior to DSS-induced intestinal injury, WAT from *Il22ra1^Adipo^* mice showed a significant reduction in *Fabp4* expression and a marked increase in the proliferation marker Ki67. These findings indicate that loss of IL-22RA1 signaling in adipocytes disrupts adipocyte differentiation and lipid metabolism, leading to increased proliferation of preadipocytes or stromal cells without proper maturation. Importantly, this altered adipose tissue response occurred despite similar levels of colonic inflammation between knockout and control mice, suggesting a critical role for adipocyte IL-22RA1 signaling in maintaining metabolic and inflammatory homeostasis in WAT during combined metabolic and intestinal inflammatory stress.

## Introduction

Interleukin-22 (IL-22), a member of the IL-10 cytokine family, is predominantly produced by immune cells such as Th17 and type 3 innate lymphoid cells (ILC3s) (1,2). IL-22 signals through a heterodimeric receptor complex composed of IL-22RA1 and the ubiquitously expressed IL-10RB (IL-10Rβ). In contrast, IL-22RA1 expression is largely restricted to non-hematopoietic cells, including epithelial cells, hepatocytes, and adipocytes (3,4). This restricted expression pattern localizes IL-22RA1 signaling primarily to barrier tissues and metabolically active organs.

Interleukin-22 (IL-22) has been shown to play an important role in intestinal defense, including regulating microbiota colonization, limiting inflammation, and promoting tissue regeneration (5). Mice lacking IL-22 or IL-22RA1 were highly susceptible to chemical and pathogen-induced colitis (6). IL-22 mounts an inflammatory response in colorectal cancer and chronic colitis models (7,8). Adipocytes express IL-22RA1, albeit at a lower level than the intestine and liver (9). Whether IL-22 confers a protective or inflammatory roles in adipocytes during intestinal inflammation remains poorly understood.

A growing body of literature has highlighted IL-22’s protective role in metabolic disorders, particularly those induced by high-fat diet (HFD) (9–11). Exogenous IL-22 administration in HFD-fed mice has been shown to improve glucose homeostasis, reduce adipose tissue inflammation, and decrease body weight (10, 12–14). In adipocytes, IL-22 has been reported to promote lipolysis by upregulating lipases such as hormone-sensitive lipase (LIPE), patatin-like phospholipase domain-containing protein 2 (PNPLA2), and fatty-acid β-oxidation (acyl-CoA oxidase 1, ACOX1), thereby lowering triglyceride content in white adipose tissue (WAT) (10). Additionally, IL-22 enhances glucose uptake in brown adipose tissue and promotes thermogenesis (10, 14). It also induces “beiging” of white fat, stimulating the formation of beige adipocytes, which are metabolically active and burn excess energy (15). While these studies demonstrate the therapeutic potential of IL-22, they rely heavily on exogenous cytokine delivery and do not isolate the direct effects of IL-22 signaling within adipocytes, especially during intestinal inflammation. To address this gap, we generated adipocyte-specific *Il22ra1* knockout mice (using the Adiponectin-Cre system).

We induced colitis in *Il22ra1^Adipo^* mice using dextran sulfate sodium (DSS) under normal chow and HFD conditions. We found significant difference in weight loss on day 9 following DSS treatment along with toward reduced colon length and decreased *Cd206* expression in distal colon tissue of *Il22ra1^Adipo^*mice fed a chow diet. Furthermore, *Il22ra1^Adipo^* mice exhibited a notable reduction in the expression of *Acox1*, a key enzyme involved in fatty acid β-oxidation, along with reduced expression of inflammation related genes in WAT. When subjected to both HFD and DSS, control mice lost substantial weight during the DSS recovery phase, while knockout mice maintained their body weight. This relative resistance to weight loss in knockouts may reflect altered adipocyte function or energy balance under combined metabolic and inflammatory stress. Importantly, *Fabp4* (fatty acid binding protein 4) expression was significantly reduced in the WAT of knockout mice. To assess adipocyte maturation, we examined the proliferation marker Ki67 and observed a marked increase in expression in knockout mice, suggesting a possible accumulation of immature or proliferating preadipocytes. Collectively, these results indicate that IL-22RA1 signaling in adipocytes plays a previously unrecognized role in maintaining proper adipocyte function, lipid metabolism, and immune interactions, particularly under conditions of metabolic and inflammatory stress.

## Materials & Methods

### Mice

Adiponectin-Cre mice on C57BL/6J background were purchased from The Jackson Laboratory. The generation of *Il22ra1^fl/fl^* mice on C57BL/6J background has been previously described (16). Mice were housed in specific pathogen free conditions at the Division of Laboratory Animal Resources at Stony Brook University. This study was approved by Stony Brook University’s Institutional Animal Care and Use Committee.

### DSS treatment

Mice (7-9 weeks old) were treated with 2.5% DSS for 8 days and then euthanized on day 10.

### DSS and HFD treatment

Mice (7-9 weeks old) were placed on HFD (Bio Serve) for 2 weeks followed by 7 days on 2.5% DSS water (MP Biomedicals). Mice were collected on day 20 from the start of DSS treatment. These mice were maintained on HFD throughout the experimental time course. Mice were separated based on genotypes before starting experiments.

### qPCR

Distal colon tissues and epididymal WAT were collected in TRIzol reagent for RNA extraction. RNA was transcribed to cDNA using a Bio-RAD iScript cDNA synthesis kit. Quantitative real-time PCR (qRT-PCR) was performed using Bio-Rad SsoAdvanced SYBR Green master mix. Primers were ordered from Integrated DNA Technologies (*Supplemental Table 1*).

### Hematoxylin &Eosin (H&E) Staining

Distal colon tissues and epididymal WAT were fixed in 10% Formalin. After paraffin embedding, 5 μm sections were deparaffinized in xylene and rehydrated with a decreasing ethanol gradient. Sections were stained with hematoxylin solution (VWR International) for 30 seconds and washed under running tap water for 15 minutes. Eosin staining was performed for 5 minutes. Slides were then dehydrated through an increasing ethanol gradient, treated with xylene, and mounted using toluene-based medium. The histological score was calculated as previously described in the chemically induced colitis model (Scheme 1) (17).

### Ki67 and Perilipin2 Immunofluorescent staining

Epididymal WAT was fixed in 10% formalin, paraffin-embedded, and sectioned into 5 µm slices. Sections were deparaffinized in xylene and rehydrated through a decreasing ethanol gradient. Antigen retrieval was performed by boiling sections in citrate buffer for 40 minutes. Tissues were then permeabilized using 0.1% Triton X-100 in phosphate-buffered saline (PBS) for 10 minutes and blocked with 5% bovine serum albumin (BSA) in PBS for 1 hour at 37 °C.

Sections were incubated overnight at 4 °C with either mouse anti-Ki-67 (1:100; BD Biosciences, Ref: 550609) or rabbit anti-Perilipin 2 (1:100; ADFP, Abcam, Ref: ab108323) primary antibodies. Sections were then washed and incubated for 1 hour at room temperature with either anti-mouse IgG (H+L) Alexa Fluor® 647 (1:100; Jackson ImmunoResearch, Ref: 715-605-150) or anti-rabbit IgG (H+L) Alexa Fluor® 488 (1:100; Jackson ImmunoResearch, Ref: 111-545-144).

Slides were washed and mounted with DAPI (Southern Biotech). Images were acquired using an Olympus CKX41 microscope. Ki67 and PLIN2 positive area percentage was calculated using Image J software.

### Statistical Analysis

Data were analyzed using GraphPad Prism version 10 software. The specific statistical tests used for each experiment are described in the corresponding figure legends.

## Results and Discussion

### IL-22RA1 signaling in adipocytes exerts a limited effect on the development of DSS-induced colitis

Our group previously demonstrated the critical role of IL-22RA1 signaling in the intestinal epithelial compartment (9, 18). We also confirmed that adipocytes express *Il22ra1*, albeit at lower levels than ileal and liver tissues (9). Given the relatively low *Il22ra1* expression in adipocytes, we hypothesized that IL-22 might have limited direct effects on adipocyte function under homeostatic conditions. To probe this pathway under inflammatory stress, we induced colitis using DSS. Notably, *Il22ra1^Adipo^*mice exhibited significant differences in colitis progression compared to littermate controls on day 9 of DSS treatment, as indicated by body weight loss. A trend toward reduced colon length was observed, while histological analysis of distal colon tissue revealed no differences in tissue damage between the two groups (*Figure 1A-C*). We measured the expression of *Il1b, Il6, Ccl2, and Cd206* in distal colon tissue to assess inflammation and macrophage polarization, as these genes are key indicators of pro-inflammatory cytokine signaling, immune cell recruitment, and anti-inflammatory macrophage activity. Analysis of inflammatory markers in the distal colon revealed no significant differences in *Il1b, Il6, or Ccl2* expression; however, we observed a trend toward reduced *Cd206* expression. Similarly, no changes were detected in genes related to lipid metabolism (*Acox1, Fabp4, Pparg*) (*Figure 1D-E*). Collectively, our data indicate that loss of IL-22RA1 signaling in adipocytes has only a minor effect on colitis development, as evidenced by modest weight reduction and a trend toward shorter colon length, without significant changes in intestinal inflammation or tissue injury in the DSS model.

**Figure 1.**
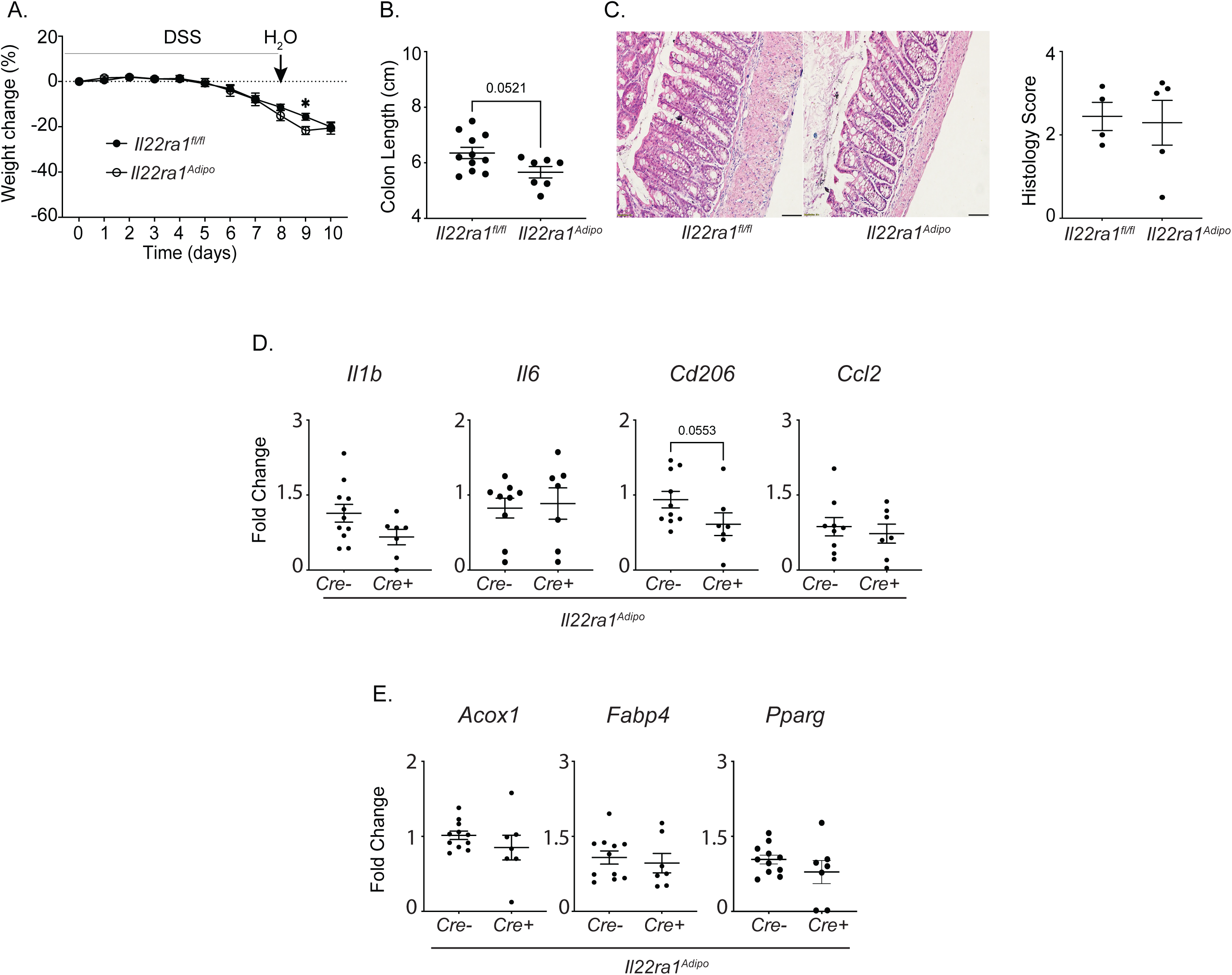
Loss of IL-22RA1 signaling in adipocytes has limited affect on susceptibility to DSS colitis. (A) Body weight change, (B) colon length, (C) H&E stained distal colon tissue with histological score in the right panel, were measured in DSS-treated *Il22ra1^fl/fl^* and *Il22ra1^Adipo^* mice. Expression of (D) immune response– and (E) lipid metabolism–related genes in distal colon tissue of DSS-treated mice was analyzed by qRT-PCR. Data are presented as mean ± SEM. Two-way ANOVA (A) and Mann–Whitney test (B–E). Data are representative of at least three independent experiments (A, B, D–E). Scale, 50 μm.

### Adipocyte-specific IL-22RA1 signaling regulates inflammatory and metabolic gene expression in WAT following colon injury

Creeping fat refers to the hypertrophy and encroachment of mesenteric adipose tissue around inflamed intestinal segments, a hallmark feature of Crohn’s disease in humans (19). Although mice can exhibit inflammation or hypertrophy of mesenteric fat, they do not develop true creeping fat as observed in patients with inflammatory bowel disease (IBD) (20). Consequently, in the present study, we focused on assessing the systemic effects of gut inflammation on visceral adipose tissue rather than direct gut–mesenteric fat interactions.

To this end, we analyzed the epididymal WAT from DSS-treated *Il22ra1*^Adipo^ mice and their littermate controls. We observed a significant reduction in epididymal WAT weight in the knockout mice compared with controls (*Figure 2A*). Consistent with this, WAT from knockout mice showed a significant decrease in inflammatory markers *Il1b* and *Cd206*, and a significant reduction in *Acox1* expression, with a trend toward decreased *Fabp4* and *Pparg* expression (*Figure 2B-C*).

**Figure 2.**
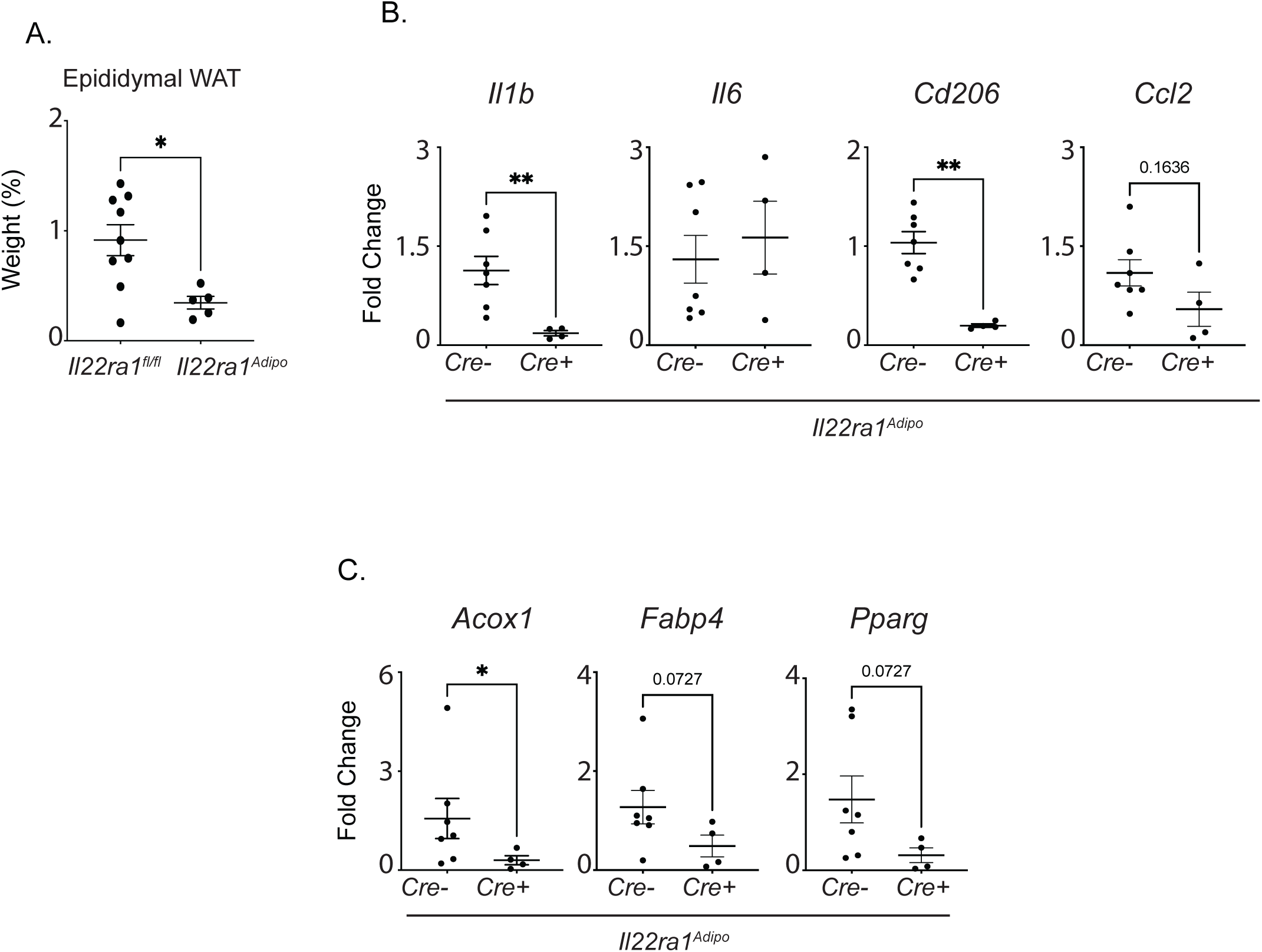
Adipocyte IL-22RA1 signaling is essential for maintaining immune balance and lipid metabolic function in WAT following colonic injury. (A) Epididymal WAT weight (% of whole body weight) of DSS-treated *Il22ra1^fl/fl^* and *Il22ra1^Adipo^* mice. Expression of (B) immune response– and (C) lipid metabolism–related genes in the WAT of DSS-treated mice were analyzed by qRT-PCR. Data are presented as mean ± SEM. \**P* < 0.05, \*\**P* < 0.01; Mann–Whitney test (A-C). Data are representative of two independent experiments (A-C).

Our previous work showed that exogenous administration of rIL-22.Fc in C57BL/6J mice did not affect adipogenesis in WAT (9). In contrast, the present study demonstrates that IL-22 released during intestinal inflammation can directly influence adipocytes. The observed reduction in WAT mass was accompanied by downregulation of genes related to inflammation and lipid metabolism, without corresponding changes in the colon. Together, these findings suggest that IL-22, produced during gut inflammation, exerts systemic metabolic effects on adipocytes via IL-22RA1 signaling, thereby linking intestinal immune responses to peripheral metabolic regulation.

IL-1β has been shown to promote adipogenesis in both murine and human adipose-derived stem cells, playing a key role in WAT remodeling by driving adipocyte differentiation (21). Moreover, IL-22RA1 signaling in WAT has been shown to induce IL-1β production (22). CD206 M2-like macrophages also contribute to WAT remodeling by promoting adipocyte progenitor proliferation and differentiation through TGF-β signaling (23). In our study, *Il22ra1^Adipo^* mice subjected to DSS-induced colitis exhibited reduced *Il1*β *and Cd206* expression in WAT, accompanied by a significant reduction in WAT weight. These results indicate that the loss of IL-22RA1 signaling in adipocytes impairs adipose tissue remodeling, likely by limiting both pro-adipogenic cytokine production and M2-like macrophage support. Consequently, IL-22RA1 deficiency may hinder adipocyte differentiation, tissue expansion, and metabolic adaptation under inflammatory stress.

### Adipocyte IL-22RA1 signaling in WAT plays a limited role in induction of DSS colitis under metabolic and inflammatory stress

Our group previously evaluated the effects of long-term HFD feeding under homeostatic conditions and found no significant differences between *Il22ra1^Adipo^* mice and controls in terms of body weight gain, glucose tolerance, WAT mass, tissue morphology, or expression of metabolic genes (9). Additionally, *Il22* transcript levels in WAT remained unchanged, suggesting that IL-22RA1 signaling in adipocytes may not be essential for maintaining metabolic homeostasis under steady-state conditions (9).

However, IL-22 production by intestinal immune cells is known to be impaired in obese mice following immune challenges, such as *Citrobacter rodentium* infection (10). Given this, and our observation that DSS-induced gut inflammation significantly altered adipose tissue in *Il22ra1^Adipo^* mice (*Figure 2*), we next subjected HFD fed mice to DSS treatment. This approach enabled us to investigate the crosstalk between WAT and the gut under combined metabolic and inflammatory stress conditions.

Following DSS treatment, littermate controls exhibited significant weight loss on days 8 and 10, while knockout mice showed less pronounced weight loss (*Figure 3A*). By day 20, both groups had recovered similarly in body weight, and no differences were observed in histological damage, indicating comparable levels of local colonic inflammation (*Figure 3A-B*). Consistent with these findings, the expression of inflammatory and lipid metabolism–related genes in the distal colon did not differ between *Il22ra1^Adipo^* and littermate control mice (*Figure 3C-D*). Together, these findings indicate that IL-22RA1 signaling in adipocytes is not required for regulating intestinal inflammatory responses, even under the metabolic stress induced by HFD feeding.

**Figure 3:**
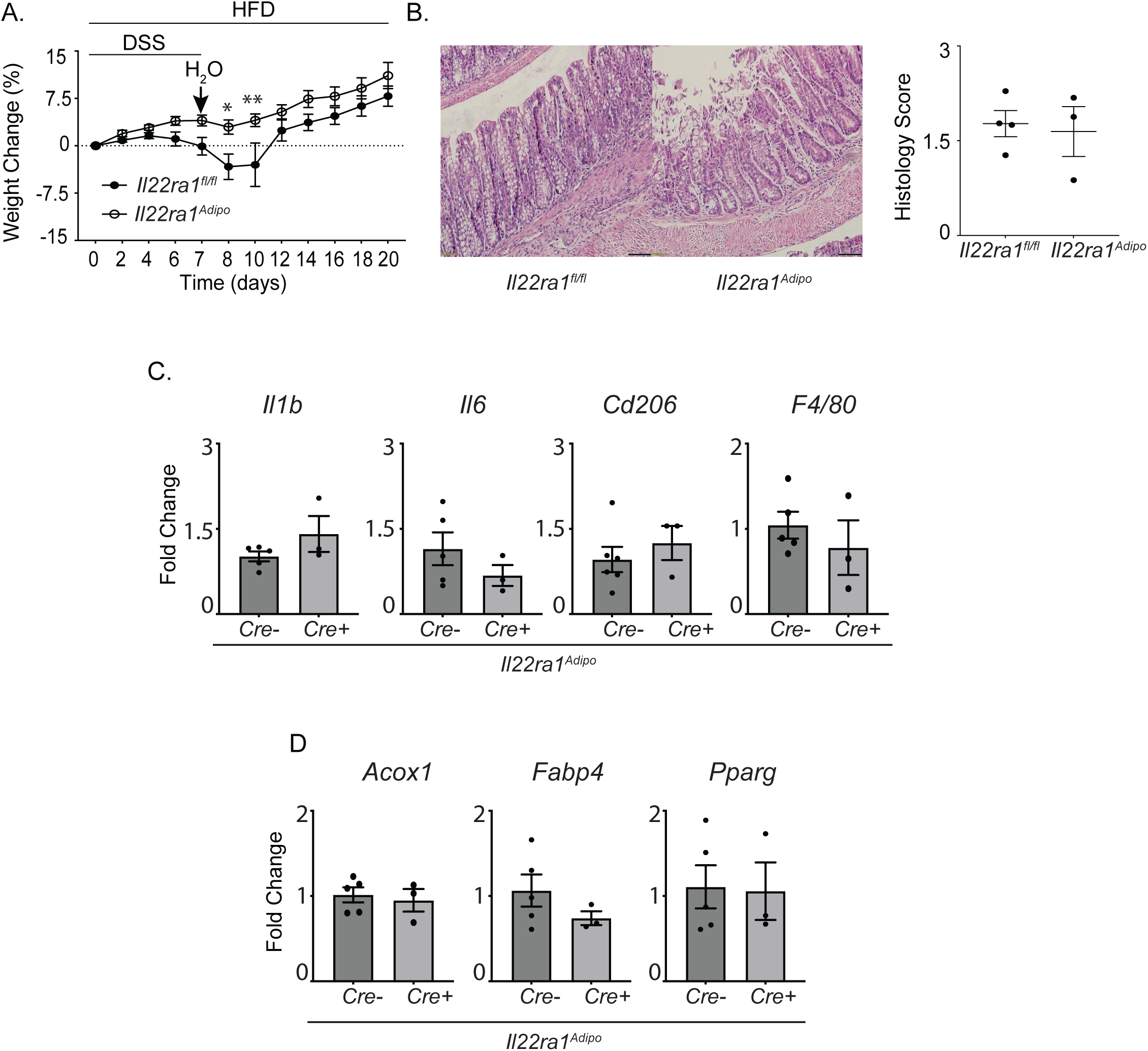
Adipocyte IL-22RA1 signaling has a modest effect on colitis development under conditions of HFD and DSS-induced inflammation. (A) Body weight change and (B) H&E stained distal colon tissue with histological score in the right panel of HFD and DSS treated *Il22ra1^fl/fl^* and *Il22ra1^Adipo^* mice. Expression of (C) immune response– and (D) lipid metabolism–related genes in distal colon tissue was analyzed by qRT-PCR. Data are presented as mean ± SEM. \**P* < 0.05, \*\**P* < 0.01; two-way ANOVA (A) and Mann–Whitney test (B-D). Results are representative of at least three (A) and two (C-D) independent experiments. Scale, 50 μm.

### IL-22RA1 activity in adipocytes sustains adipogenesis even under metabolic and inflammatory challenges

Next, we examined changes in WAT following combined HFD and DSS treatment. No differences were observed in the relative weight of epididymal WAT between groups (*Figure 4A*). However, *Il6* expression showed an increasing trend, while the macrophage-related gene *F4/80* was significantly reduced (*Figure 4B*). Moreover, the expression of the metabolic and adipocyte differentiation gene *Fabp4* was markedly decreased in knockout mice, with a trend toward reduced *Acox1* expression (*Figure 4C*). Together, these findings suggest that loss of IL-22RA1 signaling enhances adipose tissue inflammation and impairs adipocyte metabolic function under combined metabolic and inflammatory stress.

**Figure 4:**
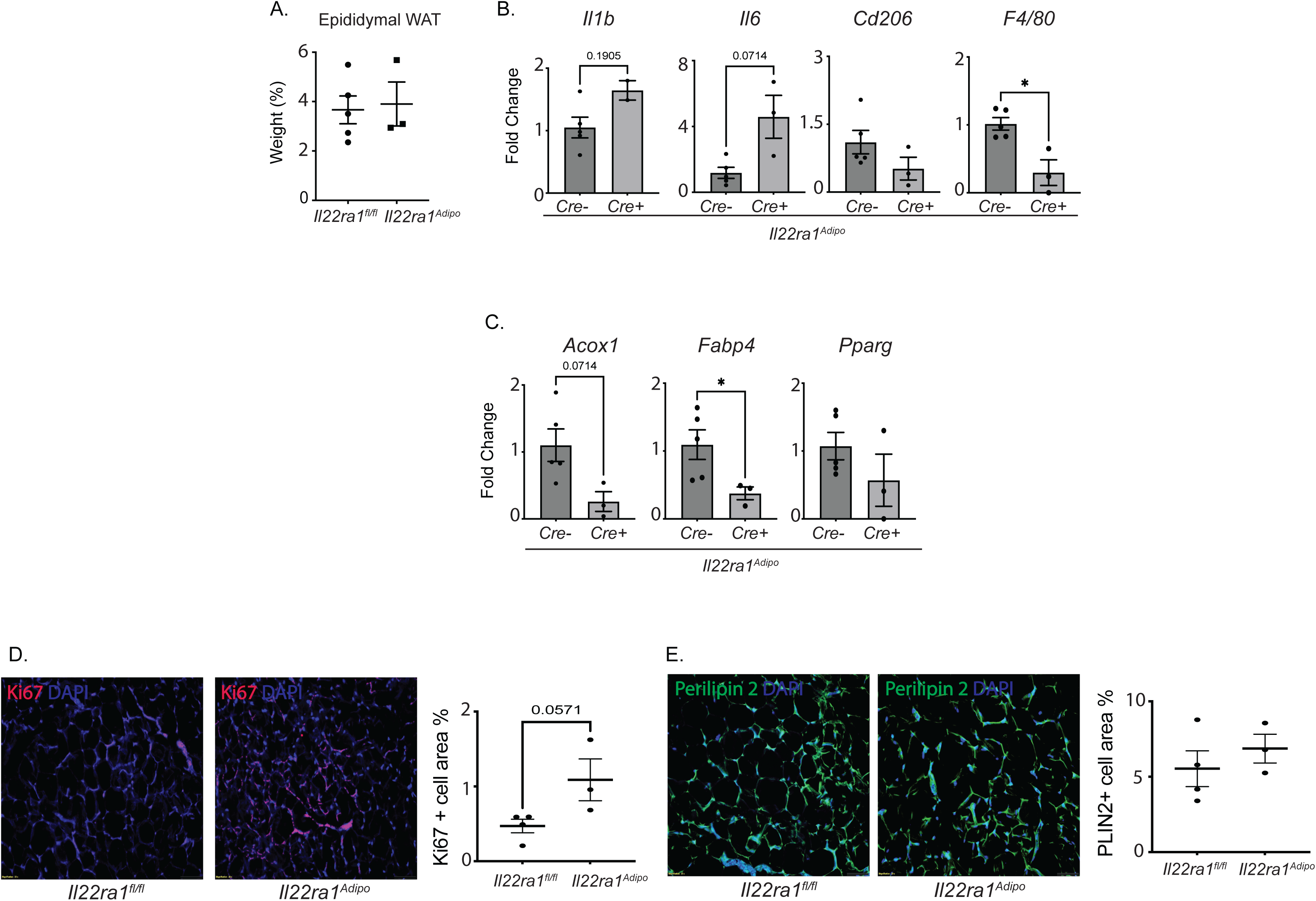
Adipocyte IL-22RA1 signaling regulates adipogenesis in WAT. (A) Epididymal WAT weight (% of whole body weight) was measured in *Il22ra1^fl/fl^* and *Il22ra1^Adipo^* mice subjected to HFD and DSS treatment. Expression of (B) immune response–related and (C) lipid metabolism–related genes in WAT from HFD- and DSS-treated mice was analyzed by qRT-PCR. Representative immunofluorescence images showing (D) Ki67 and (E) PLIN2 protein levels in WAT from *Il22ra1^fl/fl^* and *Il22ra1^Adipo^* mice under HFD and DSS treatment. Quantification of Ki67 cell area (%) and PLIN2 area (%) was performed using ImageJ software and presented in the corresponding right panels. Data are shown as mean ± SEM. \**P* < 0.05; Mann–Whitney test (A–E). Results are representative of two independent experiments. Scale bar, 50 μm.

The decrease in *F4/80* expression may reflect a reduction in macrophage numbers due to impaired recruitment or increased apoptosis, whereas the trend towards increase in *Il6* expression suggests a persistent inflammatory state, likely driven by adipocytes in *Il22ra^Adipo^* mice. Reduced macrophage presence can impact adipocyte function, as macrophages-particularly CD206 M2 macrophages-support adipocyte differentiation and metabolic activity (23). Consequently, this may contribute to the observed reduction in *Fabp4* expression.

Given the reduced expression of *Acox1* and *Fabp4*, respectively, in DSS-treated and HFD+DSS-treated *Il22ra1^Adipo^* mice, we hypothesized that adipocyte differentiation and/or lipid metabolic function might be compromised. *Acox1* actively promotes adipogenic differentiation in preadipocytes (24), while *Fabp4* serves as a well-established marker of mature adipocytes (25). Based on this, we next examined Ki67, a marker of cellular proliferation (26), and Perilipin2 (PLIN2), which is expressed during early stages of adipocyte differentiation and marks immature adipocytes (27), to assess potential disruptions in adipocyte precursor proliferation and maturation.

Measuring Ki67 and PLIN2 allowed us to determine whether the observed impairment in adipogenic gene expression was associated with (a) increased proliferation of preadipocytes or stromal cells (*Ki67* upregulation) and/or (b) accumulation of immature adipocytes (changes in PLIN2 expression). We found a marked increase in Ki67 expression, but no significant change in PLIN2, suggesting enhanced proliferation without progression toward adipocyte maturation (*Figure 4D-E*).

These findings support a model in which loss of *Il22ra1* leads to impaired preadipocyte differentiation as evidenced by reduced *Acox1* and *Fabp4* expression accompanied by a compensatory increase in cellular proliferation. However, this proliferative response appears to stall before terminal differentiation, as PLIN2 expression remains unchanged. This likely results from impaired adipogenesis and/or inflammatory remodeling driven by the absence of IL-22RA1 signaling in adipocytes.

## Conclusion

The role of IL-22RA1 signaling in adipocytes remains largely unexplored. In this study, we demonstrate the importance of IL-22RA1 signaling in adipocytes during gut inflammation. IL-22 is known to promote intestinal epithelial cell proliferation and tissue repair following injury (28). Following DSS-induced intestinal injury, our findings indicate that IL-22RA1 signaling supports adipocyte regeneration and the expression of key metabolic genes, thereby preserving adipose tissue integrity and function under inflammatory stress. Collectively, these results expand the current understanding of IL-22 biology beyond epithelial repair to include a role in inter-organ communication between the gut and adipose tissue. This newly identified function has potential implications for metabolic syndrome, IBD, and obesity-associated inflammation, conditions in which gut–adipose crosstalk is often disrupted.

## Acknowledgments

The automated tissue processing and paraffin embedding were performed by the Research Histology Core Laboratory at Stony Brook University. PK is supported by the Crohn’s and Colitis Foundation (CCF 476637 and CCF 1168040).

## Author Contributions

P.K. and A.S. conceived and designed the experiments. A.S., K.S., S.J.G., and H.T. performed the experiments. A.S. drafted the manuscript, and P.K. revised and edited it.

## Conflict of Interest Statement

The authors declare no financial or commercial conflicts of interest.

## Abbreviations

HFD: High Fat Diet
DSS: Dextran Sulfate Sodium
WAT: White Adipose Tissue
IBD: Inflammatory Bowel Disease

## Notes

### Competing Interest Statement

The authors have declared no competing interest.

