## Supplemental Table 1 for "Adipocyte IL-22RA1 signaling promotes structural and functional remodeling of white adipose tissue following acute intestinal damage"

**Supplementary Table 1:** List of mouse primers used for qPCR analysis.

| **Vendor** | **Gene** | **Primer Sequence** |
| --- | --- | --- |
| **Integrated DNA Technologies** | ***Gapdh*** | Forward: 5’-TCATCAACGGGAAGCCCATCAC-3’  Reverse: 5’-AGACTCCACGACATACTCAGCACCG-3’ |
|  | ***Acox1*** | Forward: 5’-CGCACATCTTGGATGGTAGT-3’  Reverse: 5’-GGCTTCGAGTGAGGAAGTTATAG-3’ |
|  | ***Pparg*** | Forward: 5’-AGGCGAGGGCGATCTTGACAG-3’  Reverse: 5’-AATTCGGATGGCCACCTCTTTG-3’ |
|  | ***Ccl2*** | Forward: 5'- TTA AAA ACC TGG ATC GGA ACC AA-3'  Reverse: 5'- GCA TTA GCT TCA GAT TTA CGG GT -3' |
|  | ***Il6*** | Forward: 5'- TCC AAT GCT CTC CTA ACA GAT AAG -3'  Reverse: 5'- CAA GAT GAA TTG GAT GGT CTT G -3' |
|  | ***Il1b*** | Forward: 5'- GCA ACT GTT CCT GAA CTC AAC T -3'  Reverse: 5'- ATC TTT TGG GGT CCG TCA ACT -3' |
|  | ***Cd206*** | Forward: 5'- AAG GCA TGC GTT GCA CAT AC -3'  Reverse: 5'-ATT CTG CTC GAT GTT GCC CA -3' |
|  | ***Fabp4*** | Forward: 5'- CCC GCA TGG AGG GTG TAT G-3'  Reverse: 5'- TGG AGG GAT CAC GAG CTT GAA -3' |
|  | ***F4/80*** | Forward: 5'- CTT TGG CTA TGG GCT TCC AGT C -3'  Reverse: 5'- GCA AGG AGG ACA GAG TTT ATC GTG -3' |
| **Qiagen (Cat#: QT00247667)** | ***Ki67 (Mm_MKi67_1_SG)*** | **N/A** |
